# A jump distance based parameter inference scheme for particulate trajectories in biological settings

**DOI:** 10.1101/238238

**Authors:** Rebecca Menssen, Madhav Mani

## Abstract

One type of biological data that needs more quantitative analytical tools is particulate trajectories. This type of data appears in many different contexts and across scales in biology: from the trajectory of bacteria performing chemotaxis to the mobility of ms2 spots within nuclei. Presently, most analyses performed on data of this nature has been limited to mean square displacement (MSD) analyses. While simple, MSD analysis has several pitfalls, including difficulty in selecting between competing models, handling systems with multiple distinct sub-populations, and parameter extraction from limited time-series data. Here, we provide an alternative to MSD analysis using the jump distance distribution (JDD). The JDD resolves several issues: one can select between competing models of motion, have composite models that allow for multiple populations, and have improved error bounds on parameter estimates when data is limited. A major consequence is that you can perform analyses using a fraction of the data required to get similar results using MSD analyses, thereby giving access to a larger range of temporal dynamics when the underlying stochastic process is not stationary. In this paper, we construct and validate a derivation of the JDD for different transport models, explore the dependence on dimensionality of the process, and implement a parameter estimation and model selection scheme. We demonstrate the power of this scheme through an analysis of bacterial chemotaxis data, highlighting the interpretation of results and improvements upon MSD analysis. We expect that our proposed scheme provides quantitative insights into a broad spectrum of biological phenomena requiring analysis of particulate trajectories.

## INTRODUCTION

A quantitative analysis of particulate trajectory data is increasingly common in biological studies. This type of data spans length and time scales from bacterial chemotaxis (1–6) to the motion of ms2 spots in Drosophila embryos (7). When analyzing particulate trajectories, on a basic level, one would like to know how the particles are moving (e.g. pure diffusion vs. directed-diffusion), and what parameters (e.g. diffusion constant), guide that motion. Additionally, seeing how the quantitative traits evolve over time can provide valuable insight.

### Mean squared displacement

Typically particulate trajectory analysis has been done through the use of the mean squared displacement (MSD). The most basic version of mean square displacement for N particulate trajectories is defined by Eq (1).

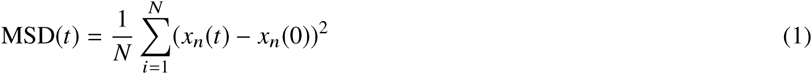

There are more complicated ways to calculate the MSD, such as doing a sweeping average (called a time-averaged MSD), but the general concept remains the same. The MSD follows well defined forms (8–10) for different modes of transport. Eq (2a)-Eq (2c) give these forms for a purely diffusive system (D), a directed diffusion system (V), and a constrained or anomalous diffusion system (A), based on how we simulated data (11).

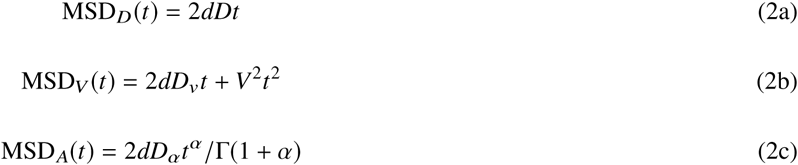

It should be noted that all diffusion can be represented by the anomalous form, but pure diffusion and directed diffusion represent simplifications for specific anomalous exponents. Additionally, for most of this paper, we will approach anomalous diffusion in the subdiffusive (*α* < 1) context, but all results do work with *α* > 1. These equations for the MSD hold across dimensions, with only a constant *d* that changes depending on the dimensionality of the system. Beyond characteristic equations, the MSD has well defined shapes when graphed, which we display in Fig. 1B for one dimensional trajectory data.

**Figure 1:**
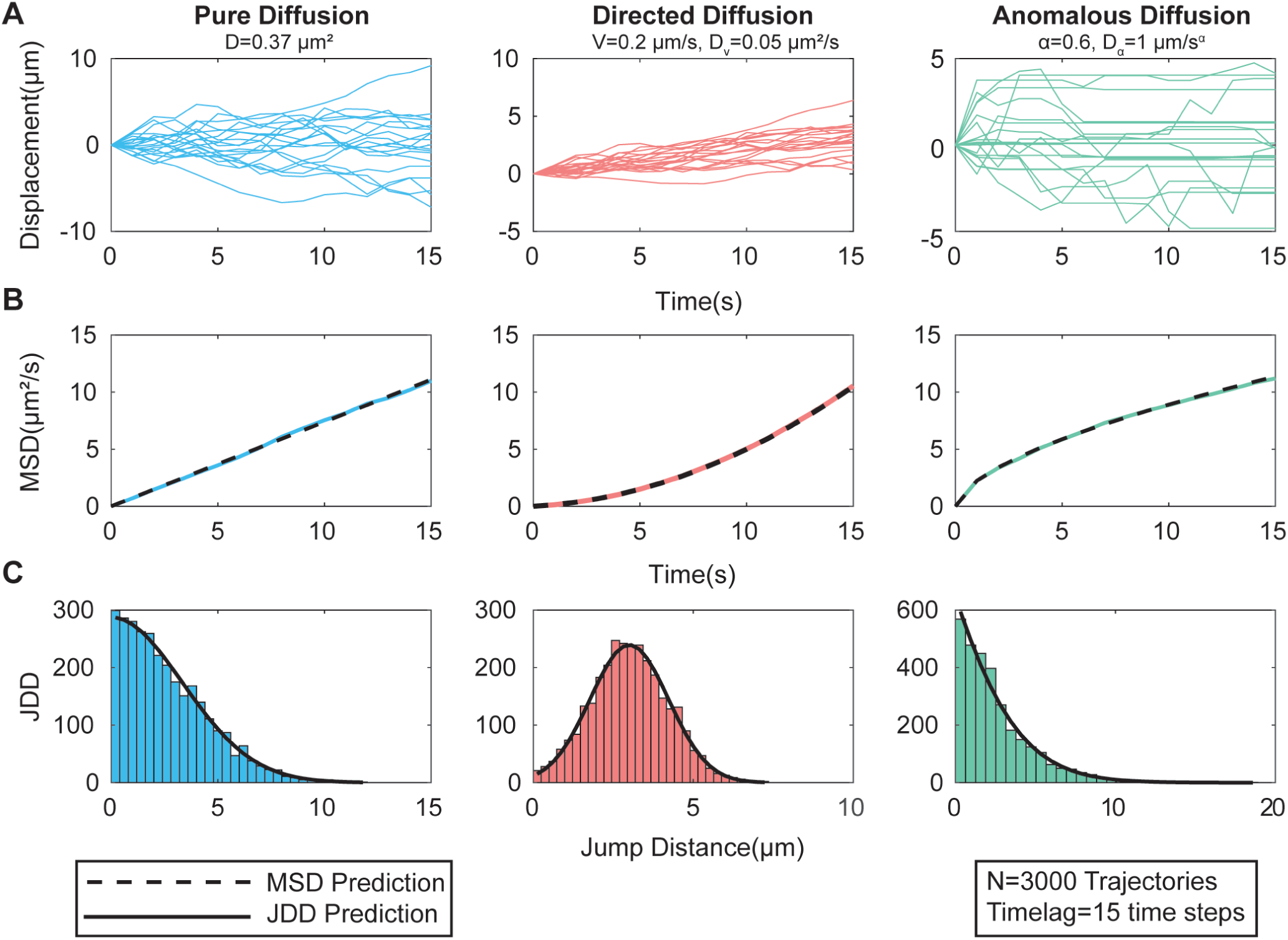
Trajectories, MSDs, and JDDs for three types of diffusion. (A) Examples of simulated trajectories are used to create MSD and JDD plots. (B) Mean squared displacement for each model and the MSD that is predicted using the simulation parameters (C) The JDD created from 3000 trajectories and the expected JDD form given the parameters of the system.

While the MSD is an established tool, it is has severe limitations. Pitfalls of MSD analysis, particularly with biological data(see Results and Discussion for examples), include the requirement for many data points (12), difficulties in selecting between competing models (13), difficulties with handling systems with two distinct subpopulations (12, 14), and the relatively large errors in cases when trajectories are short. Several studies have tried to improve and expand upon MSD analysis, and also have proposed new methods of extracting models and parameters (12–19).

### The jump distance distribution

As an alternative to the MSD, we demonstrate the power of the Jump Distance Distribution (JDD) to classify particulate trajectories (20). The JDD is closely related to the MSD. Each point on the MSD curve is the mean of the underlying JDD. Thus, with the JDD we examine a full distribution as opposed to a set of distribution means.

The idea of the JDD and its potential uses for parameter extraction is not a new one, but so far its use has been limited to purely diffusive systems in two dimensions with an assumed number of population sub-fractions (21–25) or has considered multiple models, but also only in two dimensions (20). Additionally, little work has been done on analyzing the improvement of the JDD on the MSD in anything other than two dimensional pure diffusion (24). Complete derivations of theoretical forms are missing and the treatment of directed and anomalous diffusion is lacking. This work serves to complete the fragmented picture of the JDD and serve as an easy template for model selection and parameter extraction. We also highlight the manner in which JDD results must be interpreted to shed light on experimental data.

The JDD is a frequency distribution of Euclidean distances for points separated by a time lag of τ seconds. Creating the JDD can be done by breaking up trajectory data into intervals of length τ, or by using a length τ sliding window to maximize data (see Constructing the JDD for more detailed information). Fig 1C shows example JDDs created from simulated data for three different models.

Analogous to MSD analysis, we derive the closed form mathematical solutions for the JDD (See Appendix S1 for derivations). These forms are dependent on the mode of transport and the dimensionality of the system. Table 1 lists the closed forms for pure diffusion, directed diffusion, and anomalous diffusion in one, two, and three dimensions and Fig 1C graphically shows what the closed form solution looks like upon a JDD given the simulated parameters (i.e. diffusion constant) of the system, the number of points (*N*), the time lag τ, the bin spacing (*dr*), and the bin center positions (*r*_*j*_ for bin *j*). Note that the bin spacing and bin center positions depend upon the number of bins chosen in creating the JDD frequency distribution.

**Table 1:** Closed form JDD (frequency distribution) value for bin *j* for three types of diffusion in one, two, and three dimensions

| Method | Dimension | JDD value for bin $j$ |
| --- | --- | --- |
| Pure Diffusion | 1D | $\frac{Ndr}{\sqrt{\pi D\tau}} \exp\left(\frac{-r_j^2}{4D\tau}\right)$ |
| | 2D | $\frac{Ndr r_j}{2D\tau} \exp\left(\frac{-r_j^2}{4D\tau}\right)$ |
| | 3D | $\frac{Ndr r_j^2}{2\sqrt{\pi}(D\tau)^{3/2}} \exp\left(\frac{-r_j^2}{4D\tau}\right)$ |
| Directed Diffusion | 1D | $\frac{Ndr}{\sqrt{4\pi D_V\tau}} \exp\left(\frac{-(r_j^2 + V^2\tau^2)}{4D_V\tau}\right) \left(\exp\left(\frac{Vr_j}{2D_V}\right) + \exp\left(\frac{-Vr_j}{2D_V}\right)\right)$ |
| | 2D | $\frac{Ndr r_j}{2D_V\tau} \exp\left(\frac{-(r_j^2 + V^2\tau^2)}{4D_V\tau}\right) I_0\left(\frac{Vr_j}{2D_V}\right)$ |
| | 3D | $\frac{Ndr r_j^2}{2\sqrt{\pi}(D_V\tau)^{3/2}} \exp\left(\frac{-(r_j^2 + V^2\tau^2)}{4D_V\tau}\right) \frac{2D_V \sinh\left(\frac{Vr_j}{2D_V}\right)}{Vr_j}$ |
| Anomalous Diffusion | 1D | $\frac{Ndr}{2\pi\sqrt{D_\alpha}} \int_{-\gamma+iT}^{\gamma+iT} \exp(ip\tau) (ip)^{\alpha/2-1} \exp\left(\frac{-r_j(ip)^{\alpha/2}}{\sqrt{D_\alpha}}\right) dp$ |
| | 2D | $\frac{Ndr r_j}{2\pi D_\alpha} \int_{-\gamma+iT}^{\gamma+iT} \exp(ip\tau) (ip)^{\alpha-1} K_0\left(\frac{r_j(ip)^{\alpha/2}}{\sqrt{D_\alpha}}\right) dp$ |
| | 3D | $\frac{Ndr r_j}{2\pi D_\alpha} \int_{-\gamma+iT}^{\gamma+iT} \exp(ip\tau) (ip)^{\alpha-1} \exp\left(\frac{-r_j(ip)^{\alpha/2}}{\sqrt{D_\alpha}}\right) dp$ |

The method presented here can account for two or more distinct subpopulations of motion occurring in the data, or if there is a switch in motion at a certain point, (20–23) by multiplying each type of motion by the fraction undergoing (or fraction of time in) the motion, and adding the distributions together. Our experimental results show examples of this in practice.

The paper is organized as follows. In the Methods section, we describe the JDD and its closed mathematical forms, how to simulate data and turn trajectory data into the JDD, how to approach parameter fitting, and finally how to select among competing models of motion. This gives us a complete processing pipeline that can be used to analyze particulate trajectories. In the Results and Discussion section, we show parameter fitting and model selection results for a range of simulated parameters, explain cases where the JDD provides improvement over the MSD, and present JDD analysis on bacterial chemotaxis data.

## METHODS

### Pipeline for processing data

The proposed pipeline has three major components.

1. Construct JDD
  - Collect particulate trajectory data
  - Choose lag time τ + number of bins *N*_*b*_ → construct the JDD
2. Parameter estimation
  - Use MSD to seed parameter fitting scheme + fit each model using non-linear weighted least squares → 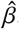, the set of Maximum-Likelihood parameters for all models.
  - Bootstrap to define error bounds on parameters → 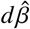.
3. Model selection
  - Integrate models over the parameter ranges 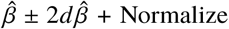 by the length of the integration range (per parameter) → *P*(*JDD*| *M*), the probability of observing the data given the model.
  - Employing Bayes Theorem gives *P*(*M* | *JDD*) → model selection.

### Creation of simulated data and JDD

#### Simulation of particulate trajectories

In this study, we use simulated data and validate our method in one dimension, but we have validated general results in two and three dimensions as well. For our simulations in one dimensions, pure diffusion was simulated using random Gaussian steps with a variance 2*Ddt* at each time point. Directed diffusion was simulated with a deterministic step of *Vdt* and a random Gaussian step of 2*D*_*V*_ *dt*. Anomalous diffusion was simulated using a continuous time random walk (CTRW) (26, 27) using a waiting time as drawn from a generalized Mittag-Leffler function (11) and a random Gaussian steps at each moving point of the size 2*D*_*α*_*dt*′^*α*^, where we set *dt*′ to be the same as the parameter *ξ* from the Mittag-Leffler function we drew from. With the CTRW, the time of each move does not correspond exactly to a set time step, requiring projection onto a predetermined grid of time steps. These simulation methods can be extended to two and three dimensions by making the Gaussian steps in each direction, and in the case of direction motion, splitting up the deterministic step into each dimension by the relevant polar and spherical transformations.

#### Constructing the JDD

Constructing a JDD requires calculating the Euclidean distances between two points on a trajectory a time lag τ apart, and binning them into a histogram with a chosen number of bins, *N*_*b*_. For simulated data, insensitivity of estimated parameters to the choice of time lag is required. Table S1 in the supporting material shows an analysis of the effect of time lag on parameter estimation for simulated data. For experimental data, there is not necessarily a “best” time lag. Instead, different time lags pull out dynamics at different time scales.

For the number of bins, as a rule of thumb, we initially choose *N*_*b*_=*N*/100, as this usually gives a sufficient number of data points for fitting without leaving empty bins. We change the number of bins as needed to improve fitting, and do not use too many bins as to have overfitting. In Table S2 in the supporting material we analyze the effect of varying the number of bins on three data sets. Too few bins and the shape of the JDD can change, greatly affecting parameter fitting by changing the skewness of the distribution. If too many bins are used, the bins can become sparsely populated (even empty), and this can have a large effect on parameter fitting accuracy. This is particularly a problem with anomalous diffusion.

Beyond the choice of a time lag and number of bins, the other choice in constructing the JDD is how much the data should overlap with itself. The naive choice in constructing the JDD from trajectory data is to split up the data into independent intervals τ/*dt* + 1 points long. While this avoids overlapping or correlated data, it requires long trajectories. As an alternative, a sliding window JDD can be employed. A sliding window can be constructed by taking a trajectory of length S and splitting it into trajectories of length τ/*dt* + 1 points long as such, [{1,1+τ/*dt*},{2,2+τ/*dt*},{3,3+τ/*dt*}…{S-τ/.*dt*-1,S}], where the numbers represent the index in a trajectory, allowing the construction of a JDD. In our initial analysis, we simulated trajectories of length τ/*dt* + 1, so every JDD data point is independent. In Table S3 and Table S4 in the supporting material, we compare non-sliding and sliding JDDs for their accuracy in parameter fitting and model selection for pure and directed diffusion.

### Parameter estimation and closed form JDDs

#### Derivation of closed form JDD equations

Our parameter estimation scheme relies on non-linear weighted least squares estimation. In order to perform this type of estimation, we require a closed form solution for each model we are examining. This required us to compile and re-derive prior work on the JDD in two dimensions (20–25), and derive the closed form solutions in one and three dimensions. In the case of pure diffusion, we can solve the relevant diffusion equation (8, 9, 22, 28). For Directed Diffusion, we were able to perform transformations on the Pure Diffusion closed form solution (8, 9). Deriving the Anomalous Diffusion form relies on finding the relevant propagator underlying an anomalous system (26). Appendix S1 in the supporting material gives full derivations for finding the JDD for each method and as a function of dimensionality. These results are compiled in Table 1.

#### Parameter estimation and estimation error

Given a constructed JDD, parameter estimation can be done through non-linear weighted least squares (NLWLS) – Eq 3. The scheme requires the sample probabilities, *y*_*j*_, and the closed form expectations, *p*_*j*_. NLWLS requires an initial guess for parameter values, which were acquired through MSD analysis of the data. Additionally, we chose to weight the scheme owing to the heteroskedasticity that is present in the errors of bin counts (See Figure S1 in the supporting material for a full explanation). We chose a weighting to be the reciprocal of the observed probabilities (1/*y*_*j*_), since our analysis demonstrates that the errors are Poissonian. To account for the possibility of empty bins, each bin was given one extra count so that no bin would be weighted with infinite weight.

To perform NLWLS on 3, we used a Levenberg-Marquardt algorithm (29, 30)on Eq 3, so in this case *β*_*i*_ is for model *i*(i being D,V, or A). We used rudimentary MSD analysis to seed the fitting for our theoretical analysis. Through our experimental analysis, we found the importance of a quality seed on fitting, so for both initial analysis and bootstrapping, we recommend using reasonable seeds, or after an initial fit, refit using the initial fit as a seed.

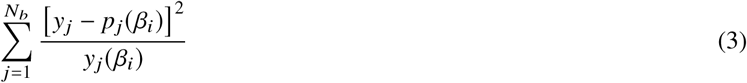

To quantify the error in our parameter fitting scheme, we employed bootstrapping (31), which helped us define an error range 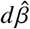, as the standard error of the bootstrapped parameters that are required in our model selection scheme.

### Model Selection

#### Bayesian inference scheme

Following the above steps we employ a Bayesian scheme to select the model that best fits the data (18, 20). The Bayesian scheme’s prior (Eq 4) assumes that all models (*M*_*i*_) are equally probable.

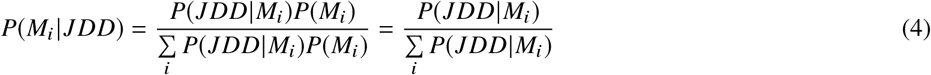

Given the fit parameters and their standard deviations, 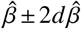 we can calculate *P*(*JDD M*_*i*_) with the probability integration scheme seen in Eq 5 (18, 20). We assume that *P*(*β*_*i*_ *M*_*i*_) is uniform based on the range of *β* and is multiplicative.

Assuming that trajectories are independent, *P*(*JDD*|*M*_*i*_, *β*_*i*_) satisfies a multinomial distribution (20), which can then be approximated by Eq 6. Even when trajectories are not independent, this approximation works well.

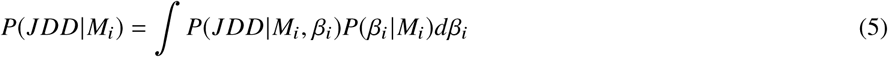

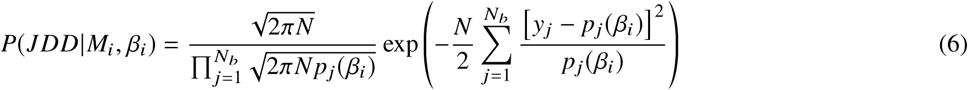

After finding *P*(*JDD, M*_*i*_) for each model, the Bayesian model selection scheme, as outlined in Eq 4, can be performed. This completes the whole pipeline, and we have posted full working examples in all three dimensions on GitHub (*github*.*com rmenssen JDD*_*Code*). We hope these examples will prove helpful on how to implement this scheme for all types of trajectory data.

## RESULTS AND DISCUSSION

We subjected the protocol outline in the methods section to the following three tests: 1) How accurate is the overall method in recovering parameter values and models, 2) what are the relative performances of the JDD and MSD based methods in cases applicable to biological data, and 3) how well does this method perform on data? What additional insights can JDD analysis provide?

Additionally, in the supporting material, we present two figures that add to our tests on the validity of the method: Figure S1, which shows the need to use weighted instead of unweighted least squares for parameter fitting, and Figure S2, which provides a broad analysis of the benefits of the JDD over the MSD across data limitations like the number of trajectories or the time lag used.

### Bayesian Inference and Parameter Estimation Results

To validate the parameter estimation and Bayesian inference scheme, we performed a broad sweep across parameters and time step to see how errors varied as a function of parameters. For each case, we performed analysis on fifty sets of trajectory data and for our error bounds bootstrapped each set fifty times to have representative parameter results and average model selection probabilities. We kept the time lag fixed at 20*dt* for the results presented in this paper, discussions of how a changing time lag affects results are left to the Supplementary Table, Table S1. We simulated 3000 independent trajectories, and used 30 bins to create our JDD. Similarly, we leave the discussion on the effect of the number of bins on parameter fitting to Table S2.

Table 2 shows average results across three different timescales for a variety of parameters. It is important to note that this table was made with JDDs that were constructed without a sliding window. Tables S3 and S4 show a comparison between non-sliding and sliding data. Non-sliding data gives more accurate results, but if data is limited, the results for using a sliding window still outperform MSD analysis.

**Table 2:**
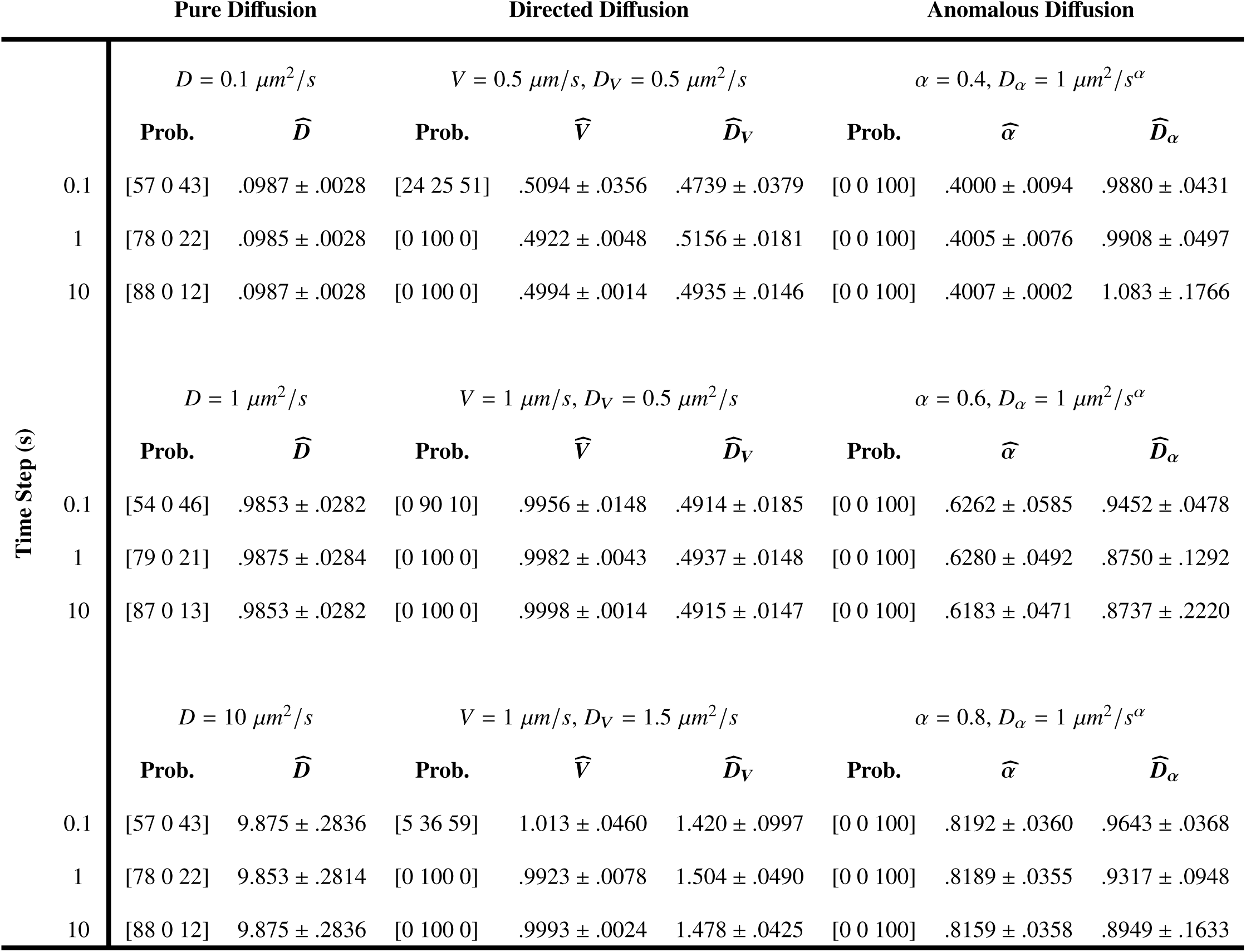
Average (50 data sets) Bayesian Selection and Parameter Estimation Results.

#### Pure Diffusion

As seen in Table 2, Pure Diffusion has the most robust results of the three models. Across the three time steps and diffusion parameters, the average error was less than two percent, and the error bound encapsulated the true simulated parameter. Often, an anomalous model with an exponent close to one is selected in preference to a purely diffusive model, which is a superficial feature of the scheme. Penalizing the number of parameters by making more complicated models less likely or by integrating over a larger parameter range suppresses this effect or after parameter fitting, excluding an anomalous model if the exponent is near one would eliminate the problem.

#### Directed Diffusion

We tested three cases of the relationship between the directed motion parameter (*V*) and the diffusion constant (*D*_*V*_): *V* ∼ *D*_*V*_, *V* > *D*_*V*_, and *D*_*V*_ > *D*, with three different time steps.

In all three cases, we had inaccuracies in model selection (but not parameter estimation) for a small time step (.1 s) that decreased with increasing its value. This can be understood straightforwardly – inaccuracies are substantial when *V* ^2^τ^2^ ∼ *D*_*V*_ τ, that is, when the length scale associated with diffusive and ballistic motion balance. An appropriate choice of τ, 5-10 times larger than τ ≫ *D*/*V* ^2^, mitigates the above inaccuracies in selection.

#### Anomalous Diffusion

Given the complexity of the functional forms of anomalous diffusion JDDs, we anticipate that parameter fitting results are less accurate in many cases. These inaccuracy stem from the multiple choices in the analysis: the number of bins, time lag, quality of seeding, and the inverse Laplace transform that is part of the closed form JDD. Numerical evaluation of the closed functional form required exploring different integration cutoffs and breaking up the domain of integration, more details of which are explored in Appendix S2 in the supporting material. The methods discussed there are possible ways to improve parameter fitting.

Regardless of the *α* used, results for fitting *α* were better than that for *D*_*α*_, with all errors on 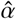 below 5% and standard errors below 10 %. Errors for *D*_*α*_ (both in terms of absolute error and standard deviation) were larger as *dt* increased. Our initial test focused on the subdiffusive regime (*α* < 1), but additional tests have shown that the method also works for superdiffusive motion (see section on Application to Chemotaxis Data).

### JDD as an improvement on MSD

We have shown that the JDD method performs well, but does it fix the pitfalls of the MSD? We demonstrate that the JDD-based method developed here is a significant improvement compared to MSD analysis, both in terms of accuracy and error bounds. We will first examine how JDD overcomes the limitations of the MSD discussed earlier: short trajectories, few data points, and combination models. These limitations often occur in biological data.

#### Improving accuracy when the MSD fails

In the case of simulations, data limitations are usually never a problem, but they are a very real issue with experimental data. MSD analysis of biological data can often fail for three reasons: 1) The trajectories are too short, 2) there are not enough trajectories, and 3) the trajectories are representing more than one mode of motion.

To best understand the improvements the JDD can bring, we outline these three cases in Fig. 2, showing comparisons of data where the MSD performs poorly, and the JDD represents a significant improvement. In this figure, the data used to create the MSD is the same data used to create the JDD. In the first two cases, we examined a directed diffusion system, which is often tricky to handle, and the last case, we look at a double diffusion system with disparate diffusion constants. For this figure, we only look at the general accuracy, not the error bounds, but as we describe later, the JDD beats the MSD on error bounds as well. With different simulations, there can be significant differences in fitting, but these are representative cases. Here, we used a naive fitting of the MSD, but even with improved fitting (such as prohibiting negative parameter values), the results do not significantly improve.

**Figure 2:**
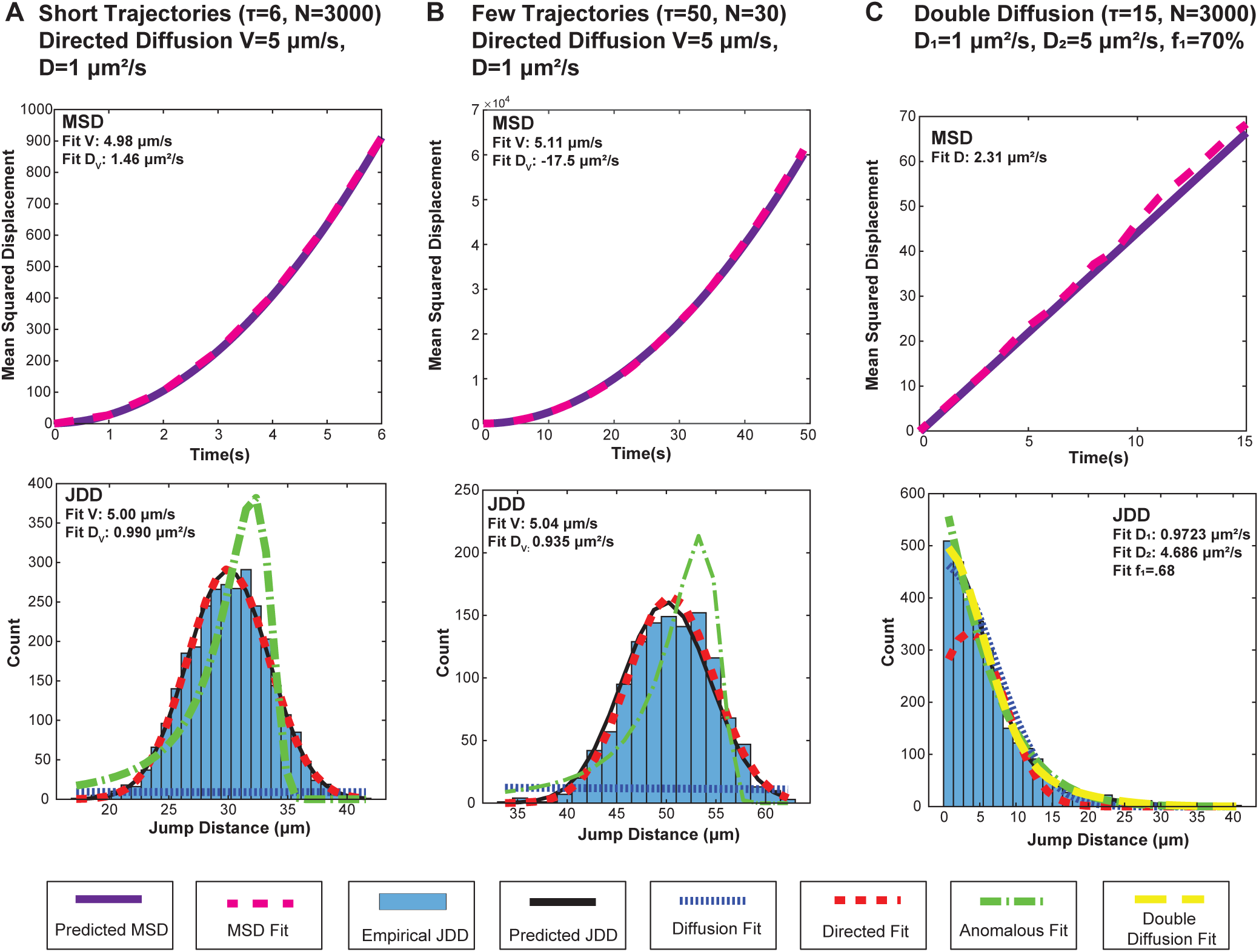
MSD vs. JDD. (A) MSD vs. JDD results in the case of short trajectories. MSD results were very inconsistent for different simulated datasets. The JDD removes this ambiguity. (B) MSD vs. JDD results in the case of few trajectories. In this case, we did not put any restrictions on the MSD fit, so the negative diffusion constant is unphysical. Even placing constraints on MSD parameters does not solve the problem and MSD fits are inconsistent from one test to the next, but JDD results are much more consistent. (C) MSD vs. JDD results in the case of two population fractions. In the MSD case, there is no way to tell that this data was created from two populations, so the MSD averages out the two. The JDD can include a double population model, and with this, it has reasonable accuracy pulling out the underlying parameters.

The first example we examine is the case where particulate trajectories are short (Fig. 2A). In our example, we simulated 3000, seven point long trajectories for directed diffusion. Examining the MSD, it is clearly directed diffusion, but doing parameter fitting on it results in almost a 50% error for this one sample (error values for other runs varied widely, another problem). This is due to the short nature of the trajectory not being able to compensate for the fact that the directed motion component is 5 times that of the diffusive component. When we perform a JDD analysis using this same data, we find great improvement in the fitting of the diffusion parameter and correct model selection. Other simulations of this system showed great variation in MSD fitting, but consistent JDD fitting.

Next, what happens in the case where particulate trajectories are long enough, but there are very few of them? (Fig. 2B). Using the same parameters and model as in the prior case, we only simulated 30 trajectories each fifty points in length. In this case, the averaging out of such a few number of trajectories produces the slightest fluctuation in the MSD curve that results in an unphysical fitting with a negative diffusion constant when using a simple fit. Even with an MSD fit banning negative parameters, the fit is still poor (*D*_*V*_ ∼ 4). This once again is an issue that is made worse in directed models over purely diffusive models.

Lastly, when examining trajectories where there are two populations (either species or behavior), the MSD is an extremely poor choice unless population subfractions are known (Fig. 2C). This happens because without knowing subfraction percentages, the MSD has infinitely many options, and the easiest is just an averaging of the different groups. So in our case, instead of getting 30% D=5, and 70% D=1, it just approximates in the middle. With the JDD, we can construct a model that allows for different population fractions, and make the naive choice that we have 50% in each population, and come out with a reasonably accurate answer. This model is then selected above the models that only assume one population, which gives it a huge advantage over the MSD.

#### MSD vs. JDD Error bound comparison

Explicit simulations showing the JDD outperforming MSD in the data-poor limit were performed, by selecting a number of trajectories, a time lag, and trajectory length and performing JDD and MSD analysis using a sliding window to construct the JDD and MSD. This analysis shows that MSD analysis has a much larger standard deviation in estimated parameter values compared to the JDD in all cases studied. Figure S2 in the supporting material is a figure showing the results of this work, and demonstrating that MSD analysis has a much larger standard deviation in estimated parameter values than the JDD in all cases studied.

### Application to Chemotaxis Data

In order to test our method as an improvement on MSD analysis, we applied our method to *E. coli* chemotaxis data (4). This data was composed of three, five minute, movies of single *E. coli* grown in a rich medium. We combined the data from the three movies to create one large data set in order to get more general results. Fig. 3A and B shows the experimental setup, as well as the three sets of trajectory data.

**Figure 3:**
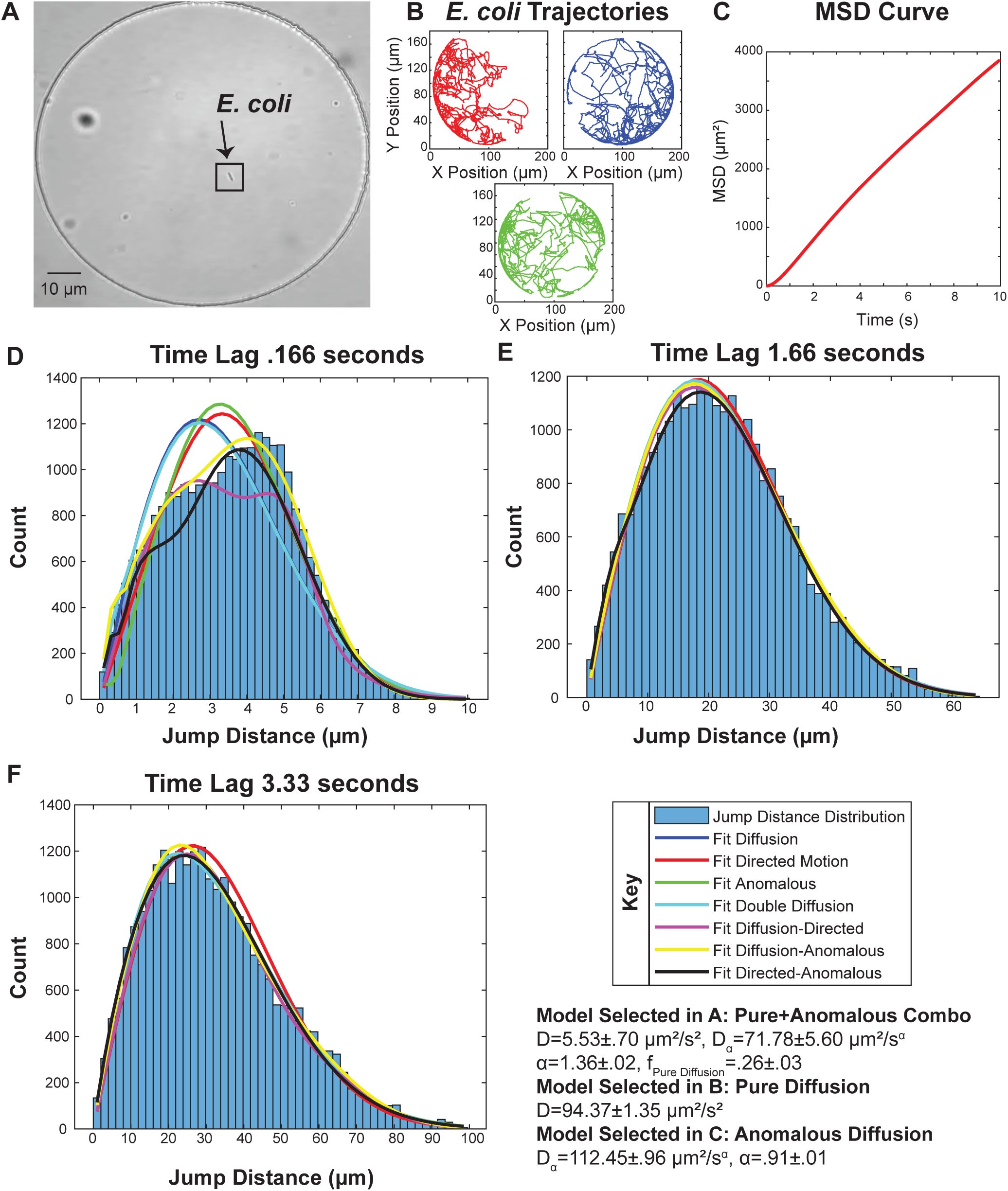
2D JDD for different time lags of Chemotaxis data. (A) Still frame of experiment with the single *E. coli* boxed in black (B) Data used in analysis. We combined all data together to make a single JDD and have a more representative data set MSD curve for up to ten seconds of time lag. Note the parabolic shape when the lag is less than 2 seconds, and how this evolves over time (D) JDD and fits for a time lag of 0.166 seconds (5 time steps) showing combination pure and superdiffusive motion as the best fit model(E) JDD and fits for a time lag of 1.66 seconds (50 time steps) showing diffusive motion as the best fit model(F) JDD fits for a time lag of 3.33 seconds (100 time steps) showing subdiffusive motion

MSD analysis resulted in poor fits and difficult to extract parameters. Additionally, the shape of the MSD curve changed from a superdiffusive to subdiffive shape, as seen in Fig. 3C. Even in the superdiffusive portion of motion, the MSD is not fit by a traditional directed diffusion model. All of these issues prompt the need for an impartial method that handles mixed models, examines of different timescales, and more accurately estimates parameters. The JDD is such a method.

Our analysis consisted of performing JDD analysis in 2D at a variety of time lags. We fit the standard models of motion (Pure, Directed, and Anomalous), as well as combination models (Double Diffusion, Pure-Directed, Pure-Anomalous, and Directed-Anomalous). Fig. 3D-F shows the result of this analysis for 3 time lags (5 steps/.166 seconds, 50 steps/1.66 seconds, and 100 steps/3.33 seconds), as well as the best fit model and its parameters.

The results of this initial analysis confirmed the presence of different dynamics at different timescales. At small timescales, the motion is a combination of pure diffusive and superdiffusive motion, being brought about by the well characterized “run and tumble” chemotactic dynamics. At further time lags, we transition to the motion appearing diffusive. This matches with the time lag at which we observe the transition from super-to-sub diffusion in the MSD analysis. At even longer timescales, the motion is subdiffusive, consistent with the averaging of run and tumble motion in a constrained environment.

Our analysis shows that with experimental data, there is no one best time lag to examine data. Different time lags will pull out different dynamics. To demonstrate this further, we performed parameter fitting over a range of time lags from 1 to 100 time steps. Fig. 4 shows the results of this analysis, in conjunction with general model fitting results for each step.

**Figure 4:**
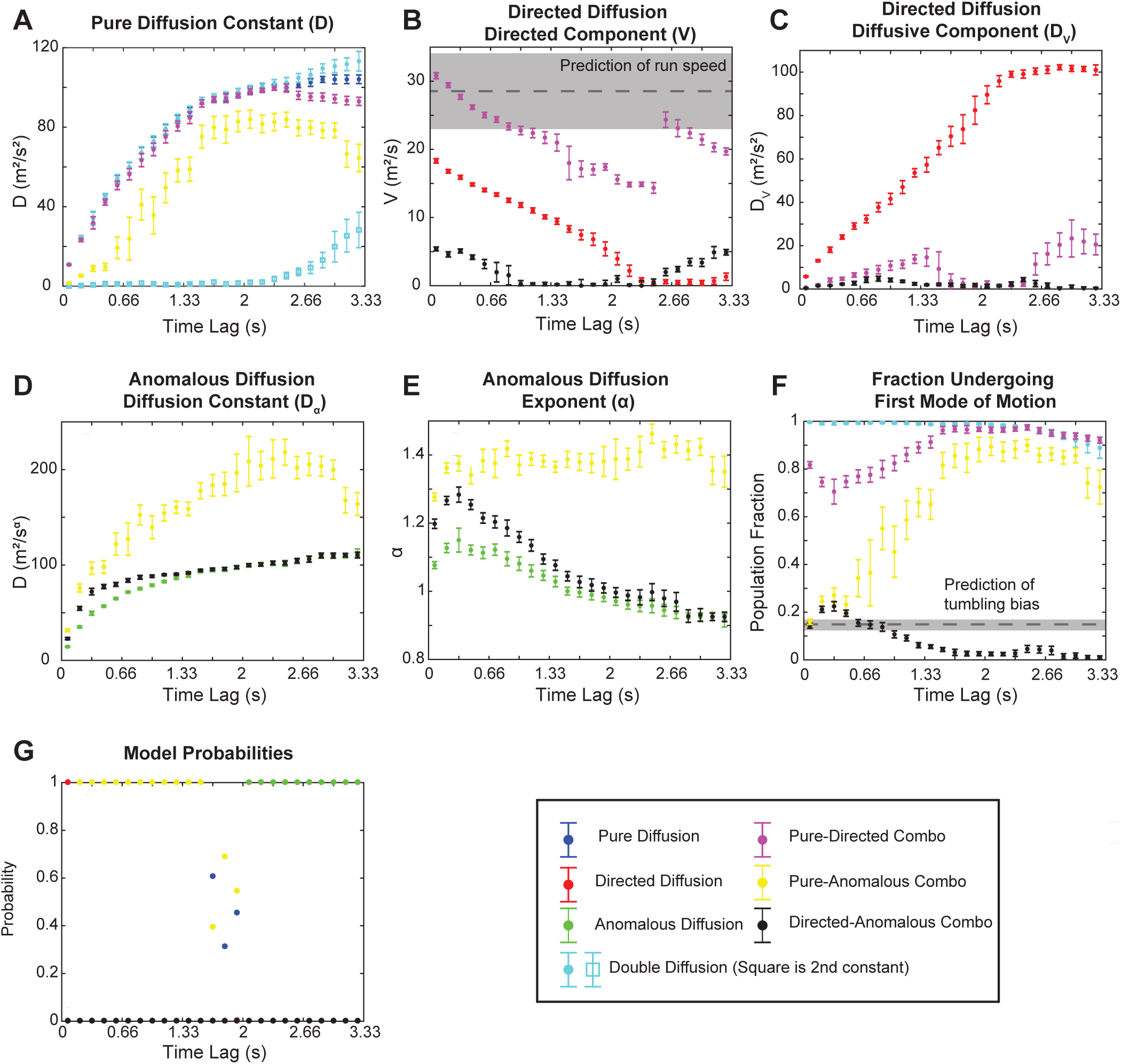
2D JDD for different time lags of Chemotaxis data. (A) All pure diffusion coefficients (*D*) (B) All directed Diffusion Coefficients (*V*). In the Pure-Directed case, the large jump around 80 time steps is due to the problem of two local minima. We compared the two options and chose the one with a lower sum of squared errors (SSE). Also, since this model was not selected, this is not a concern. At very small time lags, the Pure-Directed Model V constant provides an accurate prediction of the run speed, consistent with prior analysis on the data. The light grey box signifies the range of speeds (mean and standard deviation) noted by experiment (4). Our measured speed at short time lags is well within this range (C) All Directed Diffusion diffusion coefficients (D) All Anomalous Diffusion Coefficients (*D*_*α*_) (E) All Anomalous Diffusion Exponents (*α*) (F) All combination model first model fraction (*f*_*D*_). At small time lags, the Pure-Anomalous fraction undergoing pure diffusion provides an accurate readout of tumbling bias, or the percent of time the bacteria is tumbling, also consistent with literature. This matches with the results derived from the experimental data through tracking runs and tumbles (4). This is noted (mean and standard deviation) in grey. (G) Model selection probabilities. To derive these probabilities, we ran our model selection scheme, but eliminated models that were infeasible (for reasons such as poor fits, anomalous constants close to one, fraction under going a model near zero). At the smallest time lag (1 time step), the selection is unreliable due to noise, and this quickly disappears, and the models selection progresses as expected.

The analysis presented in Fig. 4 has shown us that in order to understand the physics at multiple timescales, we have to look at whole sweep of JDD analysis. Parameter values are slowly varying, and often asymptote, even in cases where the model is not selected. This shows power in the JDD in that we can see how dynamics change when we look at them on different scales, and can reveal real properties underlying the system. The two examples we have of this are in being able to read off the tumbling bias (fraction of time spent tumbling) and the run speed.

Our initial analysis suggested that early on, superdiffusive behavior (from running) that was mitigated to less than a pure directed model by pure diffusion (from tumbling). Our model selection early on reveals this to be true, and with a best fit model being a combination of pure and anomalous (*α*>1) diffusion. The fraction of time spent tumbling corresponds to the fraction of pure diffusion in the Pure-Anomalous model for small time steps. We get a value of.16, which is consistent with prior work done on this data (see Fig. 4F for dashed grey line representing the value and standard deviation from Fraebel et al. (4))

Despite not selecting a combination model of pure and directed motion (signifying motion of just runs and tumbles), parameter fits from this model are still interpretable. At small time steps, the speed associated with the Directed component is consistent with the measured speed given from prior analysis of the data (see Fig. 4B dashed grey line and standard deviation for values from Fraebel et al. (4)). We believe this is a unique feature one can only get from JDD analysis, and with different systems, one can expect graphs like this pulling out different underlying features of the system.

We believe this application of the JDD method to experimental data demonstrates the power of the method. We can track changes in dynamics of a system, extract a combination model (yielding an important observation about the system), and overall, our parameter estimates are greatly improved compared to the MSD fits.

## CONCLUSION

Particulate trajectory analysis is used in many different fields of study. While MSD analysis is easy to use, it often oversimplifies systems and can lead to inaccurate estimations of parameters. With increases in computing power and mathematical tools, a better and more versatile method should be used.

The aim of this paper has been to describe a general method for bringing trajectory analysis up-to-date and provide a small case study on experimental data to understand additional benefits the JDD might bring. The JDD method overcomes the issue of small amounts of data compared to the MSD, and with large amounts of data, is just as accurate. It allows for selection between competing models, which is a major advantage when uncertain of the underlying behavior of the system. It can consider combination models, which MSD analysis cannot do. The general method is broadly applicable, it works for any dimension and any model, as long as the underlying JDD frequency distribution can be derived.

We have provided the framework for implementing this model with experimental data and have posted our code online with examples for all dimensions and models. In Appendix S2 we outline tips for the application to experimental data. We also have derived and compiled the JDD frequency distributions for three modes of motion in one, two, and three dimensions. This allows for the easy creation of combination models and the ability to examine three dimensional data.

In an effort to distribute these methods widely, we have uploaded examples of each method for all dimensions for researchers to explore, and easy modifications allow for analysis of experimental data (should we put up an example experimental one?). This code is available on GitHub.

Beyond our validation of the method itself, we have shown cases where the JDD greatly outperforms the MSD, and shown an example experimental case with bacterial chemotaxis data. This case shows that JDD analysis can be thought of analyzing a family of distributions, and analyzing this family can show changing in dynamics at different time scales, and can even pull out parameters underlying the system. We hope that this analysis method can be more broadly used and bring additional accuracy and insight to researchers.

## Supporting information

## AUTHOR CONTRIBUTIONS

RM and MM designed and performed the research. RM wrote analytic tools, carried out all simulations, analyzed the data. RM and MM wrote the article.

## ACKNOWLEDGMENTS

The authors would like to thank Seppe Kuehn and Harry Mickalide at the University of Illinois at Urbana-Champaign for providing the bacterial chemotaxis data for analysis.

This material is based upon work supported by the National Science Foundation Graduate Research Fellowship Program under Grant No. DGE-1324585. Any opinions, findings, and conclusions or recommendations expressed in this material are those of the authors and do not necessarily reflect the views of the National Science Foundation

