## Supplementary material for "A jump distance based parameter inference scheme for particulate trajectories in biological settings"

### Supporting Material

### Appendix S1: Mathematical Derivations

In this section, we derive the closed form JDD expressions. We have removed any preference of direction from our distribution [1–3]. In all of our derivations, we will leave out the subscripts for bin number, use $\tau$ as our measure of time, and leave the expressions as probability distributions instead of frequency distributions (so $N$ times the distributions here will give the distributions in Table 1 in the main text).

#### Pure Diffusion

To arrive at the probability distribution describing diffusion with a constant D, one first solves the Diffusion equation for that dimension, or use a number of other means [3–6].

### 1D

From solving the basic diffusion equation, one can get a basic probability distribution for a position $r$ and time $\tau$:

$$p(r) = \frac{1}{(4\pi D t)^{\frac{1}{2}}} \exp\left(\frac{-r^2}{4D\tau}\right)$$

To turn the above expression into a jump distribution, we must account for the fact that $r > 0$, suggesting a multiplication by a factor of 2. Additionally, to generate a discrete probability distribution, we need to consider the percentage in a slice of size dr, motivating the element dr.

$$P(r) = \frac{dr}{(\pi D t)^{\frac{1}{2}}} \exp\left(\frac{-r^2}{4Dt}\right)$$

### 2D

Two dimensions proceeds similarly, by first solving the 2D diffusion equation and converting to polar coordinates.

$$p(r) = \frac{1}{4\pi D t} \exp\left(\frac{-r^2}{4Dt}\right)$$

To consider what percentage of trajectories will appear in a slice of size dr, we must integrate over the area element $rdrd\theta$, only over $\theta$ from 0 to $2\pi$.

$$P(r) = \frac{rdr}{2Dt} \exp\left(\frac{-r^2}{4Dt}\right)$$

### 3D

Solving the diffusion equation in 3D in spherical coordinates gives,

$$p(r) = \frac{1}{(4\pi D t)^{\frac{3}{2}}} \exp\left(\frac{-r^2}{4Dt}\right)$$

Integrating using spherical coordinates gives,

$$P(r) = \frac{r^2 dr}{2\sqrt{\pi}(Dt)^{\frac{3}{2}}} \exp\left(\frac{-r^2}{4Dt}\right)$$

#### Directed Diffusion

The directed diffusion closed form JDDs are a perturbation upon the pure diffusion forms. The integration follows identically, supplemented by a substitution of $r$ to $r - Vt$ must be made. Without loss of generality, we can assume that this substitution can be made along a single direction. Additionally, since all potential angles/directions of the directed motion are isotropically distributed, we must first "average" out the angular components, or in the case of one dimension, average out the two possible directions.

### 1D

In one dimension, upon substitution and averaging, we arrive at:

$$P(r)_{simp} = \frac{dr}{(4\pi Dt)^{\frac{1}{2}}} \exp\left(\frac{-(r^2 + V^2t^2)}{4Dt}\right) \exp\left(\frac{Vr}{2D}\right)$$

This is slightly incorrect though, this only considers "positive" distributions. What if the distribution would normally go negative? In this case, we need to consider wrapping those results back into the distribution. This happens mostly in the case where $V$ is small, but also broadly when $V^2t^2 \sim DV^2$, where $t = dt * timelag$

This actually means what we need is:

$$P(r) = \frac{dr}{\sqrt{4\pi D_V \tau}} \exp\left(\frac{-(r_j^2 + V^2\tau^2)}{4D_V\tau}\right) \left(\exp\left(\frac{Vr_j}{2D_V}\right) + \exp\left(\frac{-Vr_j}{2D_V}\right)\right)$$

If we are in a case where the directed component is much larger than the diffusive (or by timescales this occurs), then it simplifies to $P(r)_{simp}$, which speeds up calculations.

### 2D

For two dimensions, we start with the 2D pure diffusion $P(r)$, and make the substitutions $r = \sqrt{x^2 + y^2}$, $x \to x - Vt$, and $x = r\cos(\theta)$, gives

$$P_{temp}(r) = \frac{rdr}{2Dt} \exp\left(\frac{-(r^2 + V^2t^2)}{4Dt}\right) \exp\left(\frac{Vr\cos(\theta)}{2D}\right)$$

The last term must be averaged over to remove any angular dependence, giving us,

$$\frac{1}{2\pi} \int_0^{2\pi} \exp\left(\frac{Vr\cos(\theta)}{2D}\right) d\theta = I_0\left(\frac{Vr}{2D}\right)$$

Arriving at the finaly expression,

$$P(r) = \frac{rdr}{2Dt} \exp\left(\frac{-(r^2 + V^2t^2)}{4Dt}\right) I_0\left(\frac{Vr}{2D}\right)$$

### 3D

Starting with pure diffusion in 3D, P(r), we substitute $r = \sqrt{x^2 + y^2 + z^2}$, making the assumption that the directed motion lies along the z-axis. Thus, we choose $z \to z - Vt$, and $z = r\cos(\phi)$. Substituting, we get:

$$P_{temp}(r) = \frac{r^2 dr}{2\sqrt{\pi}(Dt)^{\frac{3}{2}}} \exp\left(\frac{-r^2}{4Dt}\right) \exp\left(\frac{Vr\cos(\phi)}{2D}\right)$$

The average has to be performed over both $\theta$ and $\phi$:

$$\frac{1}{4\pi}\int_0^{\pi}\int_0^{2\pi} \exp\left(\frac{Vr\cos(\phi)}{2D}\right)\sin\phi d\theta d\phi = \frac{\sinh\left(\frac{Vr}{2D}\right)}{\frac{Vr}{2D}}$$

Lastly, following a substitution into the original equation, we arrive at the final result.

$$P(r) = \frac{r^2 dr}{2\sqrt{\pi}(Dt)^{\frac{3}{2}}} \exp\left(\frac{-(r^2 + V^2t^2)}{4Dt}\right) \frac{\sinh\left(\frac{Vr}{2D}\right)}{\frac{Vr}{2D}}$$

#### Anomalous Diffusion

The method used to derive the following results comes from The Random Walk's Guide to Anomalous Diffusion [7] from the method of propagators. While closed form solutions do exist, they are extremely complicated, suggesting a numerical treatment of the inverse Laplace Transforms.

### 1D

We first must find the relevant propagator, $W$ [7]. Given the probability density function (pdf) for the waiting time and the pdf for the distances jumped (with variance $\sigma^2$), we perform the following manipulations. First, we convert the waiting time and jump-distance distributions to their Laplace and Fourier Transforms, respectively. For anomalous diffusion in 1D, this gives in time and space:

$$w(u) = 1 - (u\tau)^{\alpha}$$
$$\lambda(k) = 1 - \sigma^2 k^2$$

Given these two transforms we can use the following form to get the form of the propagator.

$$W(k,u) = \frac{1 - w(u)}{u}\frac{W_0(k)}{1 - \lambda(k)w(u)}$$

In this case, $W_0(k)$ is the Fourier Transform of the initial condition, which is assumed to be the Dirac Delta function.
Substituting in $w(u)$ and $\lambda(k)$, we arrive at,

$$W(k,u) = \frac{[W_0(k)/u]}{1 + D_\alpha u^{-\alpha}k^2}$$

Taking the Inverse Fourier Transform, gives us

$$W(r,u) = \frac{1}{2\sqrt{D_\alpha}} u^{\alpha/2-1} \exp\left(-ru^{\alpha/2}/\sqrt{D_\alpha}\right)$$

What remains here is taking the inverse Laplace transform. To perform an inverse Laplace transform we substitute $u$ by $ip$, since this form works better for the computational integral. Multiplying the equation by a factor of 2 and $dr$ as we have done before, brings us to the final form.

$$\frac{dr}{2\pi\sqrt{D_\alpha}} \int_{-\infty}^{\infty} \exp\left(ip\tau\right)(ip)^{\alpha/2-1} \exp\left(-r(ip)^{\alpha/2}/\sqrt{D_\alpha}\right) dp$$

##### 2D and 3D

The 2D result was recovered, after correction, from [1]. The 3D result followed straightforwardly from the 1D and 2D results.

### Appendix S2: Application to Experimental Data

Applying the JDD method to experimental data requires some additional considerations compared to simulated data. We have compiled here a few tips for ways to approach using this method in practice in order to get the most accurate results.

#### Data creation and comparison

When creating the JDD, be aware of any gaps in the data as to not have points that are multiple time points apart next to each other. The same advice holds for the basic MSD analysis used in seeding for the NLWL – always account for missing data. Otherwise, the seeds used might be closer to a different local (and not global) minimum, which will likely produce poor parameter fitting.

As evidenced in our supplemental sections, the choice in time lag, bin size, or sliding vs. non-sliding can affect the accuracy of parameter fitting results. Thus, we suggest creating a general use code that can quickly create JDDs based on these types of things to check the stationarity and accuracy of results under different conditions of creation of the JDD.

#### Stationarity of results

If comparison across different time lags or bin sizes is not stationary, there are a few potential causes beyond too little data or too short of a time lag (a particular problem if directed motion is the correct model).

One option for a non-stationary system is that there are multiple models underlying the system. Multiple models can occur if there are two distinct subpopulations in the data. With multiple modes of motion and only analyzing from a single motion perspective, changing time lags or bin sizes might change which mode is being selected, or just give inconclusive results. To check for this, including a combination models in analysis can extract those underlying modes.

The appearance of something like multiple modes without truly being two subpopulations of data can appear if data covers a time range long enough that the dynamics of the system have changed.

If when analyzing data this might be the case, then attempting a combination model and then looking at fractional splits of data based on that, or just splitting your data at a point where another important factor (like fluorescence) abruptly changes dynamics can help separate data into sections that can be analyzed individually.

#### Anisotropy

Since we have presented results for one, two, and three dimensions, we suggest analyzing data in one dimension, and then comparing to two or three dimensions to check for anisotropy. Otherwise, an important system behavior is lost, and parameters in the higher dimension will be an average of the true dynamics.

#### Parameter fitting tips

##### Anomalous Diffusion

Anomalous diffusion parameter fitting is the most difficult of the modes explored in this study. This is due to the implementation of the inverse Laplace transform numerically. In Matlab, we used the built in integral function. With this, we also had to select tolerances, integration cutoffs, and whether to split the integral into pieces or not. There were difficulties with numerical accuracy, and we do not expect these difficulties to disappear with experimental data.

A first step in trying to improve parameter fitting is to change relative and absolute error tolerances within the fitting scheme since this can create large changes in accuracy. Tracking the sum of squared errors between a JDD created by fit parameters compared to the experimental JDD is helpful in determining the magnitude of any changes.

A second parameter that can be changed to improve fittings is the integration cutoff parameter $\gamma$. In our work, we found that a smaller $\alpha$ required a larger integration range. So for example, when we were simulating data with $\alpha = 0.4$, we used a range of ˜1250 to 1250, compared to a range of -500 to 500 for $\alpha = 0.6$ and 0.8.

Occasionally, it can be advantageous to split up the anomalous integral into multiple parts. In Matlab, the integral function can only lay down a certain number of points, and the anomalous integral is highly oscillatory, so breaking the integral into smaller subintervals can allow each region to have enough integration waypoints to capture the oscillation correctly. In our work, it was necessary to do this is 3D to get accurate parameter fitting results, and was occasionally helpful in 1D.

**Is the fit bad or is the model just wrong?**

When parameter fitting is not working as it should, it can be difficult to tell if the parameter fitting is the problem, or is this the best the fitting can do because the model is just incorrect? The way to approach this problem depends on the dimension of the system. In two and three dimensions, the JDDs for each model created from the fit parameters can all essentially be the same, but in fact they come from three very different underlying behaviors. In this case, the parameter fitting is working as it should, since the end goal should be three models that fit the data. In this case, the model selection step will help discern the correct model, but if the model selected seems incorrect based on other information about the system, we suggest examining the probability integration scheme for what might be occurring.

In one dimension, the fit parameter JDDs for each model are rarely similar. At this point, it is difficult to tell if the fit truly is bad, or if it is the best that can be done. For example, when simulating pure and anomalous diffusion, directed diffusion cannot seem to fit the JDD at all. So, in these cases, the model itself is the problem, not the fit.

If there are reasons to think the issue is not the model but the fit, there are a few things that can be done to improve the fit. For pure and directed diffusion, a few error tolerances can be changed, but more likely, improved fits will be found by changing time lags or bin sizes. For anomalous diffusion, these things still hold true, but there are also the suggestions in the above section that might improve the fit.

#### Supporting Figures

**Figure S1**: Typically, when one uses weights with least squares methods, the weights are proportional to the variance of the data. If the distribution within each bin is Poissonian, then we expect the variance ((Predicted Counts-Actual Counts)$^2$) to be equal to the mean (Actual Counts).We confirm these with simulations. For each method, we simulated 500 JDDs, and computed the average value of, $[N * y_j - N * p_j(\beta_M)]^2$. This represents the variance in the data, which scales linearly as a function of the average count per bin. Our results for all three transport modes are shown below. The distribution is manifestly Poissonian, justifying our weighting scheme.

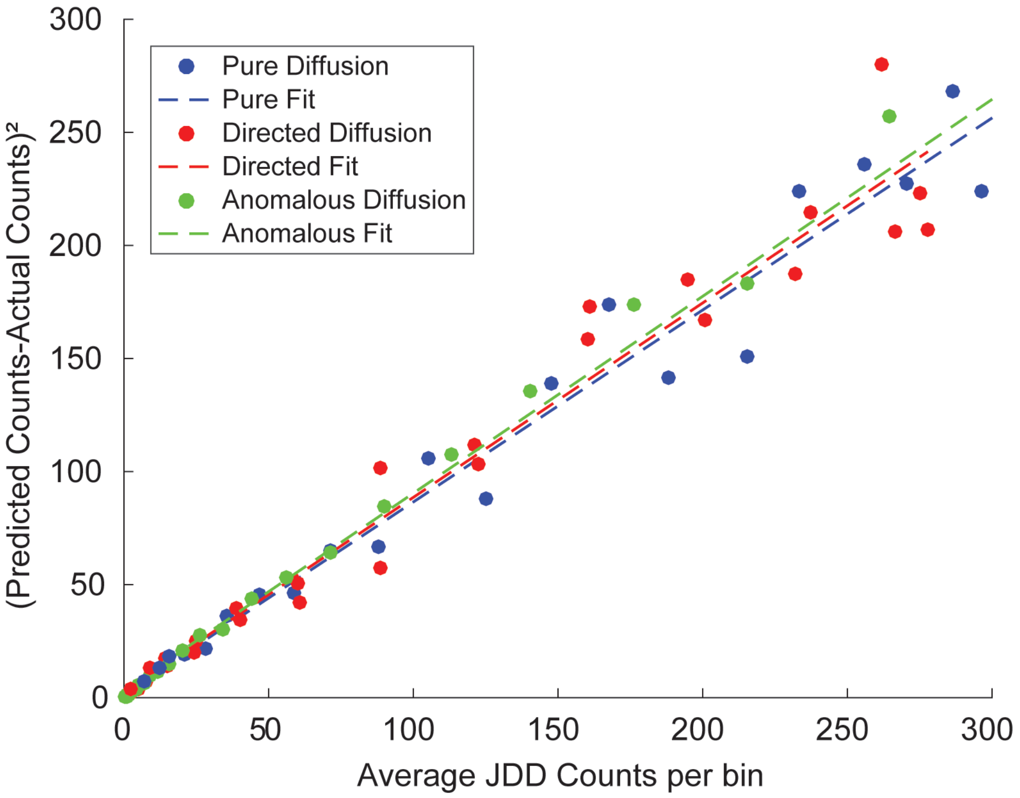


Figure 1: **Error Model for the Jump Distance** The squared error of the predicted JDD counts and the actual JDD counts compared to the actual JDD counts. The linear relationship between the two suggests that the errors are Poissonian in nature, and thus we should use a weighting of $1/y$, where $y$ is the actual JDD probabilities, to implement in our weighted least squares fitting.

**Figure S2:** For this figure, we analyzed the standard deviation of parameter values by using bootstrapping(which we use to define our error bounds) for both MSD and JDD. MSD analysis has a much larger standard deviation in estimated parameter values than the JDD in all cases studied. The JDD continues to improve upon the MSD as we have longer trajectories, more trajectories, and a shorter time lag, all of which lead to more data points. Relative performance of the two schemes was stable across parameter regimes explored.

With directed motion, it reflects that there is a time lag "sweet-spot" for best results (For more on this see Table S2). MSD analysis performs particularly poorly when the drift term ($V$) is significantly larger than the diffusion term ($D_V$), as was the case in our analysis. In this case, MSD analysis cannot reliably extract the diffusion parameter. This leads us to another large advantage of JDD analysis; when the directed part of motion is much larger than the diffusive part, JDD analysis can reliably extract the diffusion constant, whereas MSD cannot.

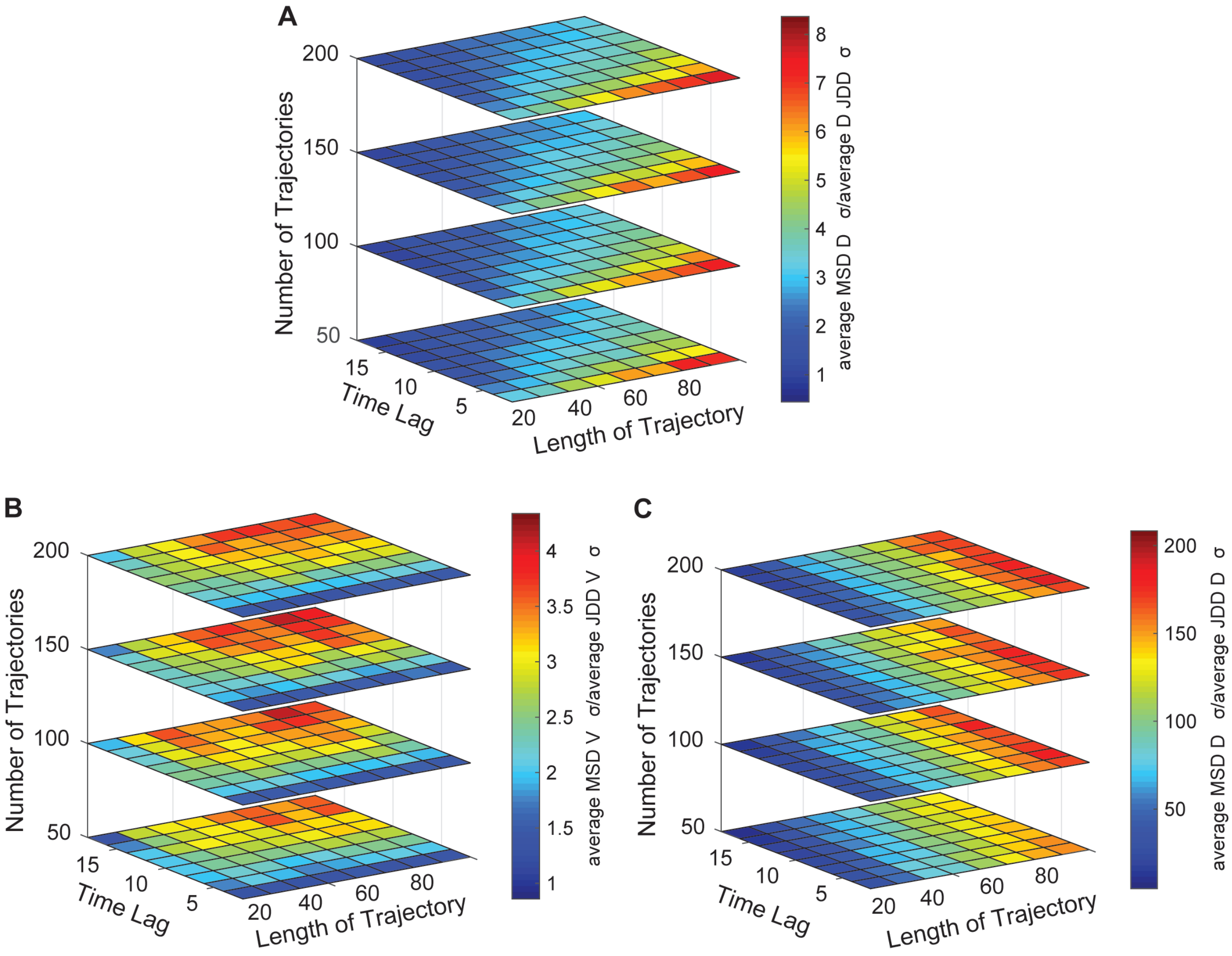


Figure 2: **MSD vs. JDD** (A) Average MSD $\sigma$/average JDD $\sigma$ for pure diffusion: D=1 $\mu m^2/s$ and dt=1 $s$ (B) Average MSD $\sigma$/average JDD $\sigma$ for directed diffusion V parameter: V=1 $m/s$ , D=0.1 $\mu m^2/s$ and dt=1 $s$. (C) Average MSD $\sigma$/average JDD $\sigma$ for directed diffusion $D$ parameter: V=1 $m/s$, D=0.1 $\mu m^2/s$ and dt=1 $s$.

#### Supporting Tables

**Table S1:** This table shows how parameter fitting results respond to different time lags. The JDD was constructed with 3000 independent trajectories of length time lag + 1, 30 bins, and a time step of 1 second. Each set was bootstrapped 50 times and refit to get the standard deviation of the fit parameters. It was then repeated 20 times and the median parameter value and standard deviation was taken for the table.

For diffusion, as we increase the time lag, the fitting improves. With a larger time lag, we should see wider variation in jump distance, which means the JDD will be filled out closer to the underlying distribution, so this result is as we expect.

With a short time lag, directed motion parameter fitting is poor. This comes from the interplay between the diffusion and directed motion constants and the overall time $\tau$, and as expected as we increase the time lag, the results improve.

For anomalous diffusion, time lag in this case does not seem to have a large effect on the parameter fitting results. This might be for a few different reasons, likely the bin size and integration cutoff (see **Appendix S2** for a discussion of this). A more complete study varying these two things is needed to show any major effects.

Table 1: **The Effect of time lag on parameter estimation**

| | $D = 1\ \mu m^2/s$ | $V = 1\ \mu m/s,\ D_V = 1\ \mu m^2/s$ | | $\alpha = .8\ \mu m/s,\ D_\alpha = 1\ \mu m^2/s$ | |
|---|---|---|---|---|---|
| **Time Lag** | $\widehat{\boldsymbol{D}}$ | $\widehat{\boldsymbol{V}}$ | $\widehat{\boldsymbol{D}}_{\boldsymbol{V}}$ | $\widehat{\boldsymbol{\alpha}}$ | $\widehat{\boldsymbol{D}}_{\boldsymbol{\alpha}}$ |
| 5 | $.9794 \pm .0283$ | $.9944 \pm .0122$ | $.9920 \pm .0352$ | $.8160 \pm .0364$ | $.9352 \pm .0613$ |
| 10 | $.9893 \pm .0294$ | $.9974 \pm .0089$ | $.9893 \pm .0299$ | $.8240 \pm .0355$ | $.9444 \pm .0757$ |
| 15 | $.9846 \pm .0276$ | $.9934 \pm .0071$ | $.9991 \pm .0299$ | $.8152 \pm .0368$ | $.9358 \pm .0945$ |
| 20 | $.9931 \pm .0302$ | $.9995 \pm .0063$ | $1.004 \pm .0302$ | $.8200 \pm .0349$ | $.9246 \pm .0915$ |
| 25 | $.9915 \pm .0292$ | $.9983 \pm .0055$ | $.9944 \pm .0303$ | $.8136 \pm .0371$ | $.9343 \pm .1065$ |

**Table S2:** This table shows how the number of bins affects parameter estimation. The JDD was constructed with 3000 independent trajectories of length 21, and with a time step of 1 second. Each set was bootstrapped 50 times and refit to get the standard deviation of the fit parameters. It was then repeated 20 times and the median parameter value and standard deviation was taken for the table.

In general, as we increase the number of bins, the error in the fit parameter increases. The standard deviation is relatively stable, since that appears to be the general accuracy for these sets of parameters. Thus, the systematic error is increasing as we increase the number of bins. This can possibly be offset by changing the time lag or in the case of anomalous diffusion, changing the integration limits.

Table 2: **The Effect of number of bins on parameter estimation**

| | $D = 1\ \mu m^2/s$ | $V = 1\ \mu m/s,\ D_V = 1\ \mu m^2/s$ | | $\alpha = .8\ \mu m/s,\ D_\alpha = 1\ \mu m^2/s$ | |
|---|---|---|---|---|---|
| **Number of bins** | $\widehat{\boldsymbol{D}}$ | $\widehat{\boldsymbol{V}}$ | $\widehat{\boldsymbol{D}}_{\boldsymbol{V}}$ | $\widehat{\boldsymbol{\alpha}}$ | $\widehat{\boldsymbol{D}}_{\boldsymbol{\alpha}}$ |
| 10 | $1.016 \pm .0289$ | $.9996 \pm .0063$ | $1.048 \pm .0308$ | $.8119 \pm .0363$ | $.9855 \pm .1045$ |
| 30 | $.9931 \pm .0302$ | $.9995 \pm .0063$ | $1.004 \pm .0302$ | $.8200 \pm .0349$ | $.9246 \pm .0915$ |
| 50 | $.9775 \pm .0301$ | $.9987 \pm .0066$ | $.9753 \pm .0302$ | $.8239 \pm .0358$ | $.9197 \pm .0929$ |
| 70 | $.9692 \pm .0300$ | $.9984 \pm .0065$ | $.9718 \pm .0305$ | $.8377 \pm .0362$ | $.8726 \pm .0927$ |
| 90 | $.9589 \pm .0305$ | $.9987 \pm .0068$ | $.9624 \pm .0304$ | $.8315 \pm .0367$ | $.8776 \pm .0865$ |

**Table S3:** We wanted to compare the sliding and non-sliding methods for making the JDD, both in model selection and parameter fitting accuracy. To do this we chose a time lag of 20 and we wanted 3000 data points composing the JDD. This implied that for the non-sliding JDD that we used 3000 trajectories, and then for the sliding, we chose to use 300, 30 point trajectories. We performed our regular analysis on the resultant JDDs(bootstrapping 50 times to get standard deviation of fit parameters, and overall repetition of 50 trials and taking of medians).

In general, the probability that the simulated data was correctly selected as diffusion was lower for sliding data than non-sliding data. The errors for each method are comparable, but we did find that if you took the standard deviation of the 50 data sets and not the median of the standard deviations compiled through bootstrapping, that this error is twice as large. This suggests that the errors listed here as the standard deviations show the inherent error in the method, and not necessarily the additional error that results from a sliding window JDD. A user should always check how these compare to errors in a larger simulated set without bootstrapping.

Table 3: **Pure Diffusion non-sliding vs. sliding Bayesian Selection and Parameter Estimation Results**

| Time Step (s) | **Non-Sliding** 3000 21 point trajectories | | **Sliding** 300 30 point trajectories | |
|---|---|---|---|---|
| | $D = 0.1\ \mu m^2/s$ | | | |
| | **Probabilities** | $\widehat{\boldsymbol{D}}$ | **Probabilities** | $\widehat{\boldsymbol{D}}$ |
| 0.1 | [56.55 0 43.45] | $.0987 \pm .0028$ | [49.69 0 50.31] | $.0962 \pm .0028$ |
| 1 | [78.33 0 21.67] | $.0985 \pm .0028$ | [68.33 0 31.67] | $.0962 \pm .0030$ |
| 10 | [87.86 0 12.14] | $.0987 \pm .0028$ | [80.38 0 19.62] | $.0692 \pm .0028$ |
| | $D = 1\ \mu m^2/s$ | | | |
| | **Probabilities** | $\widehat{\boldsymbol{D}}$ | **Probabilities** | $\widehat{\boldsymbol{D}}$ |
| 0.1 | [53.89 0 46.11] | $.9853 \pm .0282$ | [48.07 0 51.93] | $.9618 \pm .0299$ |
| 1 | [79.21 0 20.79] | $.9875 \pm .0284$ | [68.60 0 31.40] | $.9617 \pm .0280$ |
| 10 | [86.97 0 13.03] | $.9853 \pm .0282$ | [80.82 0 19.18] | $.9618 \pm .0299$ |
| | $D = 10\ \mu m^2/s$ | | | |
| | **Prob.** | $\widehat{\boldsymbol{D}}$ | **Prob.** | $\widehat{\boldsymbol{D}}$ |
| 0.1 | [56.55 0 43.45] | $9.875 \pm .2836$ | [49.73 0 50.23] | $9.617 \pm .2802$ |
| 1 | [78.35 0 21.65] | $9.853 \pm .2814$ | [68.28 0 31.72] | $9.618 \pm .2990$ |
| 10 | [88.00 0 12.00] | $9.875 \pm .2836$ | [80.36 0 19.64] | $9.617 \pm .2802$ |

**Table S4:** Using the same methods as **Table S3**, we analyzed non-sliding JDDs vs. sliding JDDs for directed diffusion data. Once again, the probabilities were lower for sliding, but given how quickly the method gives us 100% probabilities (only an issue with small time steps, which directed diffusion already struggles with), it wasn't a concern. In general, parameter fitting was slightly worse for sliding data. Once again, care must be used in setting error bounds.

Table 4: **Directed Diffusion non-sliding vs. sliding Bayesian Selection and Parameter Estimation Results**

| | Non-Sliding 3000 21 point trajectories | | | Sliding 300 30 point trajectories | | |
|---|---|---|---|---|---|---|
| | $V = 0.5\ \mu m/s$, $D_v = .5\ \mu m^2/s$ | | | | | |
| Time Step (s) | **Prob.** | $\widehat{V}$ | $\widehat{D}_V$ | **Prob.** | $\widehat{V}$ | $\widehat{D}_V$ |
| 0.1 | [24 25 51] | $.5094 \pm .0356$ | $.4739 \pm .0379$ | [16 37 47] | $.5172 \pm .0323$ | $.4606 \pm .0353$ |
| 1 | [0 100 0] | $.4922 \pm .0048$ | $.5156 \pm .0181$ | [ 0 99 1] | $.4929 \pm .0046$ | $.5086 \pm .0176$ |
| 10 | [0 100 0] | $.4994 \pm .0014$ | $.4935 \pm .0146$ | [0 100 0] | $.4997 \pm .0014$ | $.4808 \pm .0153$ |
| | $V = 1\ \mu m/s$, $D_v = .5\ \mu m^2/s$ | | | | | |
| | **Prob.** | $\widehat{V}$ | $\widehat{D}_V$ | **Prob.** | $\widehat{V}$ | $\widehat{D}_V$ |
| 0.1 | [0 90 10] | $.9956 \pm .0148$ | $.4914 \pm .0185$ | [0 71 29] | $.9978 \pm .0144$ | $.4887 \pm .0192$ |
| 1 | [0 100 0] | $.9982 \pm .0043$ | $.4937 \pm .0148$ | [0 100 0] | $.9998 \pm .0044$ | $.4777 \pm .0155$ |
| 10 | [0 100 0] | $.9998 \pm .0014$ | $.4915 \pm .0147$ | [0 100 0] | $.9997 \pm .0014$ | $.4794 \pm .0147$ |
| | $V = 1\ \mu m/s$, $D_v = 1.5\ \mu m^2/s$ | | | | | |
| | **Prob.** | $\widehat{V}$ | $\widehat{D}_V$ | **Prob.** | $\widehat{V}$ | $\widehat{D}_V$ |
| 0.1 | [5 36 59] | $1.0132 \pm .0460$ | $1.4204 \pm .0997$ | [11 47 42] | $1.017 \pm .0437$ | $1.378 \pm .0938$ |
| 1 | [0 100 0] | $.9923 \pm .0078$ | $1.504 \pm .0490$ | [0 100 0] | $.9937 \pm .0078$ | $1.472 \pm .0504$ |
| 10 | [0 100 0] | $.9993 \pm .0024$ | $1.478 \pm .0425$ | [0 100 0] | $.9997 \pm .0024$ | $1.447 \pm .0465$ |
